# A novel epitope tagging system to visualize and monitor antigens in live cells with chromobodies

**DOI:** 10.1101/2020.06.08.140327

**Authors:** Bjoern Traenkle, Sören Segan, Philipp D. Kaiser, Ulrich Rothbauer

## Abstract

Epitope tagging is a versatile approach to study different proteins using a well-defined and established methodology. To date, most epitope tags such as myc, HA, V5 and FLAG tags are recognized by antibodies which limits their application to fixed cells, tissues or protein samples. Here we introduce a broadly applicable tagging strategy utilizing a short peptide tag/chromobody (PepTag/PepCB) system. The addition of the small PepTag does not interfere with the examined structures in different cellular compartments and its detection with the fluorescently labeled Pep-chromobody (PepCB) enables optical antigen tracing in real time. By employing the phenomenon of antigen-mediated chromobody stabilization (AMCBS) using a turnover-accelerated PepCB we demonstrated that the system is suitable to visualize and quantify changes in Pep-tagged antigen concentration by quantitative live-cell imaging. We expect that this novel tagging strategy offers new opportunities to study the dynamic regulation of proteins, e.g. during cellular signaling, cell differentiation, or upon drug action.

## Introduction

Antibodies are still the key players to detect and study given proteins in biomedical research. However, their large size and multi-domain nature as well as their instability in the intracellular milieu limits their application within living cells or organisms. Single-domain antibody fragments (referred to as nanobodies, Nbs) derived from heavy-chain-only antibodies of camelids have become an attractive alternative to conventional antibodies ^1,2^. Due to their natural single-domain structure, small size, high stability and solubility, Nbs have beneficial properties for intracellular applications and there are numerous reports available which describe the selection of nanobodies for live-cell studies ^3–8^. To visualize and monitor endogenous antigens in their native surroundings, Nbs are genetically fused to fluorescent proteins to generate so-called chromobodies (CBs) which are introduced as DNA-encoded expression constructs in living cells ^9^. During the last decade multiple CBs have been developed and successfully applied in various approaches to monitor dynamic changes of their antigens in cell models and organisms (reviewed in ^2^). Despite their potential to directly probe native antigens, the *de novo* generation of gene-specific CBs is still cumbersome and time-consuming ^10–15^. Thus, generic CBs recognizing protein- or peptide-tags would be a promising alternative. A well-established example is the GFP-CB which has become widely used for multiple functional and imaging applications ranging from targeted relocalization ^9,16,17^ induced proteasomal degradation ^18,19^, or high-throughput translocation assays ^20^ of GFP-tagged proteins. However, it has to be considered that the addition of large protein tags such as GFP can affect expression, cellular localization, folding and/or function of the fusion protein ^21,22^. Thus, CBs targeting smaller peptide-derived epitope tags would be favorable. Notably, most Nbs bind conformational epitopes which strongly limits the identification of peptide specific Nbs ^23–25^. To date only very few peptide-binding Nbs are reported ^26–28^. Recently, Strokappe et. al. described a Nb (VHH 2E7) which binds with high affinity to a helix-forming peptide located in the first heptad repeat of the glycoprotein 41 (gp41) of HIV ^29^. Considering the viral origin of the peptide and the lack of homologous sequences within proteins of eukaryotic cells (blast.ncbi.nlm.nih.gov/Blast.cgi), we reasoned that the isolated peptide would provide an interesting epitope tag which could be matched by a VHH 2E7-derived CB. For a better understanding we refer to the helical epitope sequence (AVERYLKDQQLLGIW) as “PepTag” and the VHH 2E7 as “PepNb” or “PepCB” respectively. In this study, we demonstrate the functionality and application of the novel PepNb and PepCB for biochemical and live-cell studies. We show that the PepCB is capable of visualizing the dynamic distribution of Pep-tagged proteins in living cells. Additionally, following our recent developments in generating turnover-accelerated chromobodies, which can be used to optically monitor dynamic changes in protein concentration in live cells ^30^, we demonstrate that a turnover-accelerated PepCB is stabilized in the presence of its antigen. Based on our findings, we propose that the PepTag/PepCB pair yields a versatile system to use a small generic epitope tag in combination with a specific chromobody for multiple live cell studies.

## Material and Methods

### Expression constructs

Oligonucleotide sequences used for cloning are listed in **Table S1**. Expression constructs and cell lines used in this study are listed in **Table S2**. For bacterial expression of PepNb, coding sequence of the PepNb was produced by gene synthesis (Integrated DNA Technologies, Iowa, USA) and cloned into the pHEN6 vector ^31^, thereby adding a C-terminal His_6_-Tag for immobilized metal affinity chromatography (IMAC) purification as described previously ^32^. For generating Pep-chromobody (PepCB) expression constructs either comprising eGFP or TagRFP as fluorescent moiety, the PepNb sequence was ligated into BglII- and BstEII-digested backbone plasmids previously described as lamin-chromobody (eGFP) ^7^ and PCNA-chromobody (TagRFP) ^33^ (kindly provided by ChromoTek GmbH, Planegg-Martinsried, Germany). To generate homologous DNA for AAVS1 PepCB integration, first, eGFP of the previously described donor plasmid EF1-α-Ub-R-ACT-CB ^34^ was replaced by TagRFP using BamHI and XbaI restriction sites. Secondly, the ACT-Nb was replaced by PepNb using PstI and BspEI. The expression construct coding for C-terminally Pep-tagged mCherry-vimentin (mCherry-VIM_Pep_) was generated by site-directed mutagenesis of mCherry-vimentin ^14^ using the primer set *vimentin-PepTag-for* and *vimentin-PepTag-rev*. The amplified DNA was purified, digested by the restriction enzyme KpnI, and religated. Accordingly, the N-terminally Pep-tagged eGFP (_Pep_GFP) construct was produced from BC2T-eGFP ^27^ using the primer pair *PepTag-eGFP-for* and *PepTag-eGFP-rev*. Utilizing the primer set *PepTag-ACTB-for* and *PepTag-ACTB-rev* _Pep_actin was generated from BC2T-actin ^35^. The expression construct coding for _Pep_GFP-tubulin was generated by replacing PAmCherry in the plasmid PAmCherry-tubulin (PAmCherry-a-tubulin was a gift from Vladislav Verkhusha; addgene plasmid #31930) using NheI and BsrGI. For the GFP-PCNA_Pep_ expression construct a C-terminal PepTag was introduced by site-directed mutagenesis of the previously described GFP-PCNA ^36^ using the primers *PCNA-PepTag-for* and *PCNA-PepTag-rev*. To generate the expression construct for _Pep_GFP-NLS the NLS sequence was introduced at the C-terminus to the above described PepTag-eGFP with primers *eGFP-NLS-fwd* and *eGFP-NLS-rev*. The expression construct coding for eGFP-NLS was produced from pEGFP-N1 (Takara Bio USA, Inc., Mountain View, CA, USA) accordingly. All generated expression constructs were sequence analyzed after cloning.

### Cell culture, transfection, CRISPR and compound treatment

HEK293T and U2OS cell lines were obtained from ATCC (CRL3216, HTB-96), and transgenic BHK cells containing multiple *lac* operator repeats from T. Tsukamato ^37^ (Cold Spring Harbor University, New York, NY, USA). The cell lines were tested negative for mycoplasma using the PCR mycoplasma kit Venor GeM Classic (Minerva Biolabs, Berlin, Germany) and the Taq DNA polymerase (Minerva Biolabs). Since this study does not include cell line-specific analysis, cell lines were used without additional authentication. Cell lines were cultured according to standard protocols. Briefly, growth media containing DMEM (high glucose, pyruvate, ThermoFisher Scientific, Waltham, MA, USA) supplemented with 10% (v/v) fetal calf serum (FCS, ThermoFisher Scientific), L-glutamine (ThermoFisher Scientific) and penicillin/streptomycin (ThermoFisher Scientific) was used for cultivation. Cells were routinely passaged using 0.05% trypsin-EDTA (ThermoFisher Scientific) and were cultivated at 37°C in a humidified chamber with a 5% CO2 atmosphere. Plasmid DNA was transfected with Lipofectamine 2000 (Thermo Fisher Scientific) in U2OS cells according to the manufacturer’s instructions. Polyethylenimine (PEI, Sigma-Aldrich, St. Louis, MO, USA) was used for plasmid DNA transfection of HEK 293T and transgenic BHK cells as previously described ^8,27^.

For site-directed integration of the PepCB into AAVS1 genomic locus, 5 x 10^5^ U2OS cells were co-transfected with 4.5 μg of the respective donor plasmid and 0.5 μg plasmid expression vector coding for Cas9 nuclease and gRNA specific for the AAVS1 locus. 24 h post transfection cells were subjected to a 48 h selection period using 1 μg/mL puromycin dihydrochloride (Sigma-Aldrich). Puromycin-resistant cells were expanded for one week before single clones were derived from the cell pool by limiting dilution. To verify site-directed integration of the CB-donor plasmid at the AAVS1 locus, genomic DNA of individual clones and the respective parental cell line was isolated using QIAamp DNA mini Kit (QIAGEN, Hilden, Germany) according to manufacturer’s instructions. Next, primer pair AAVS-1-vor-HA-L-for and AAVS1-T2A-rev (AAVS1-primer-pair-1) primer set, and AAVS1-HA-L-for and AAVS1-HA-R-rev (AAVS1-primer-pair-2) were used for PCR-based genotyping (strategy outlined in **Supplementary Data 2**). Successful CB integration into the AAVS1 locus results in an amplicon of 1018 bps using AAVS1-primer-pair-1. To determine whether a homozygous or a heterozygous CRISPR event occurred AAVS-1-primer-pair-2 was utilized: homozygous CRISPR events result in an amplicon of 4346 bps, while heterozygous CRISPR events result in two amplicons of 4346 bps and 289 bps. Compound treatment with 2 μM cytochalasin D (Sigma-Aldrich) was performed for 10 min, followed by an exchange to cytochalasin D-free medium for additional 35 min.

### Recombinant protein production and nanobody labeling

PepNb comprising a C-terminal Sortase-tag was expressed, purified and site-directed conjugated to Alexa Fluor 647 (AF647) or ATTO488-coupled peptides H-Gly-Gly-Gly-Doa-Lys-NH2 (sortase substrate, Intavis AG, Köln, Germany) as previously described ^35,38^. Briefly, 25 μM nanobody, 75 μM dye-labeled peptide dissolved in sortase buffer (50 mM Tris, pH 7.5, and 150 mM NaCl) and 100 μM sortase were mixed in coupling buffer (50 mM Tris, pH 7.5, 150 mM NaCl, and 10 mM CaCl_2_) and incubated for 5 h at 25 °C. Uncoupled nanobody and sortase were depleted using Ni-NTA resin (Biorad, Hercules, CA, USA). Unbound dye was removed using Zeba Spin Desalting Columns (ThermoFisher Scientific). The dye-labeled protein fraction was analyzed by SDS-PAGE followed by fluorescent scanning on a Typhoon Trio (GE-Healthcare, Chicago, IL, USA, excitation 633 nm, emission filter settings 670 nm BP 30) and subsequent Coomassie staining. For sortase-coupled nanobodies DOLs of 0.7□±□ü.15 were determined.

### Immunoprecipitation, SDS-PAGE and Western Blot

3 × 10^6^ HEK 293T cells were seeded in 100 mm culture dishes (Corning, New York, NY, USA) and cultivated for 24 h. In the following, cells were subjected to plasmid DNA transfection with equal amounts of expression vectors. Subsequently, cells were washed and harvested in PBS, snap-frozen in liquid nitrogen and stored at −20 °C. Cell pellets were homogenized in 100 μL lysis buffer (10 mM Tris/HCl, 150 mM NaCl, 0.5 mM EDTA, 0.5% NP40, 1 mM PMSF, 1 μg/mL DnaseI, 2.5 mM MgCl_2_, 1 x protease inhibitor cocktail (Serva, Heidelberg, Germany)) by repeated pipetting and vortexing for 60 min on ice. Lysates were clarified by subsequent centrifugation at 18,000 x g for 15 min at 4 °C. The supernatant was adjusted with dilution buffer (10□mM Tris/HCl, 150 mM NaCl, 0.5 mM EDTA, 1 mM PMSF) to 0.5 mL. 10 μL (2%) were added to 2 x SDS-containing sample buffer (60 mM Tris/HCl, 2% (w/v) SDS, 5% (v/v) 2-mercaptoethanol, 10% (v/v) glycerol, 0.02% bromphenole blue; referred to as input). For immunoprecipitation purified PepNb was immobilized to NHS-activated sepharose (GE-Healthcare) as described previously ^15^. 30 μL of sepharose-coupled nanobodies were added to the protein solution and incubated for 16 h on an end-over-end rotor at 4 °C. After centrifugation (2 min, 2500 g, 4 °C) supernatant was removed and the bead pellet was washed two times in 0.5 mL dilution buffer, resuspended in 2 x SDS-containing sample buffer and boiled for 10 min at 95 °C. Denaturing polyacrylamid gel electrophoresis (SDS-PAGE) was performed according to standard procedures. For western blotting proteins were transferred on nitrocellulose membrane (GE Healthcare).

### Intracellular Immunoprecipitation (IC-IP)

3 × 10^6^ HEK293T cells were transiently co-transfected with equal amounts of expression vectors coding for PepCB (TagRFP) and _Pep_GFP or eGFP. 24 h after transfection cells were harvested, lysed as described and PepCB or eGFP were precipitated using the RFP-trap or GFP-trap (ChromoTek) according to manufacturer’s protocol. Input and bound fractions were subjected to SDS-PAGE followed by immunoblotting analysis with anti-TagRFP antibody (Evrogen, Moskau, Russia) and anti-GFP antibody (3H9, ChromoTek).

### Immunofluorescence

For immunofluorescence staining, U2OS cells were plated at 8 x 10^3^ per well in a μClear 96 well plates (Greiner Bio-One, Kremsmünster, Austria). Next day, cells were transfected with plasmids coding for _Pep_actin or GFP-PCNA_Pep_. 24 h after transfection, cells were washed twice with PBS and fixed with 4% (w/v) paraformaldehyde (PFA) in PBS for 15 min at RT or with a 1:1 mixture of methanol/acetone for 15 min at −20 °C. Incubation with labeled PepNbs (50 ng/mL in 5% BSA in TBST) was performed overnight at 4 °C. For nuclear staining 4’,6-diamidino-2-phenylindole (DAPI, Sigma-Aldrich) was used. Unbound nanobody and DAPI were removed by three washing steps with PBS and images were acquired immediately afterwards with a MetaXpress Micro XL system (Molecular Devices, San Jose, CA, USA).

### Microscopy and FRAP analysis

For fluorescence recovery after photobleaching (FRAP) experiments, 8 x 10^3^ U2OS cells per well were seeded in a Cellstar 96-well plate (Greiner Bio-One) and transiently transfected with cDNAs encoding GFP-actin, actin-CB or PepCB in combination with and _Pep_actin. FRAP recordings were performed with a Zeiss confocal laser scanning microscope (CLSM 510 Meta) using a 488 nm argon laser and 63 × magnification. For photobleaching, the laser was set to 20% output and 100% transmission to bleach a 2.2 × 2.2 μm region of interest for 2.4 s. Confocal imaging series were acquired with 1% laser transmission and the pinhole opened to 1.5 Airy units. Generally, 5 prebleach and 145 postbleach images were recorded with 588 ms time intervals. Normalized mean fluorescence intensities were corrected for background and for total loss of fluorescence over time. Fluorescence recovery curves were fitted with Origin 7.5 using an exponential function, given by *I*(*t*) = *A*(1 − *e^-kt^*), where *I(t)* is the signal intensity dependent on time, *A* is the end value of intensity, *k* is the time constant. Half-times of recovery were determined by 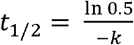.

### Image Segmentation and Analysis

8 x 10^3^ cells were plated per well in a black μClear 96-well plate (Greiner Bio-One) and transfected as described above. For automated nuclear segmentation, nuclei of live cells were stained by addition of 2 μg/mL Hoechst33258 (Sigma-Aldrich) to the cell culture medium. Images were acquired with an ImageXpress micro XL system (Molecular Devices) and analyzed by MetaXpress software (64 bit, 6.2.3.733, Molecular Devices). Fluorescence images comprising a statistically relevant number of cells were acquired for each condition (>200 cells). For quantitative fluorescence analysis the mean fluorescence of the PepCB in cells transiently expressing a PepTag-comprising antigen or its respective control was determined. Custom Module Editor (version 2.5.13.3) of the MetaXpress software was used to establish an image segmentation algorithm based on the parameters size and fluorescence intensity above local background. An image segmentation mask was generated based on the PepTag-comprising antigen or its control. The average chromobody fluorescence intensities were determined for each image followed by subtraction of background fluorescence. From these values the mean fluorescence was determined and standard errors were calculated for three independent biological replicates and student’s t-test was used for statistical analysis.

## Results and discussion

### Generation of a PepTag/PepNb capture system

To test whether the PepNb is able to capture recombinant proteins comprising a PepTag, we covalently coupled purified PepNb to sepharose beads to generate an affinity matrix that we refer to as Pep-nanotrap. For immunoprecipitation we incubated soluble protein lysates of HEK293T cells transiently expressing N-terminally Pep-tagged eGFP (_Pep_GFP) or eGFP as control (**Figure 1 A**) with the Pep-nanotrap and analyzed the input, non-bound and bound fractions by SDS-PAGE followed by Coomassie staining and immunoblotting. Pep-nanotrap specifically precipitated _Pep_GFP but not eGFP (**Figure 1 B**). However, immunoblotting revealed that a significant amount of _Pep_GFP remained in the non-bound fraction indicating that the Pep-nanotrap was not able to quantitatively deplete _Pep_GFP from the cellular lysate (**Figure 1 B**). This might be due to a high dissociation rate of the antigen which is supported by an improved pulldown efficiency when using a greater amount of Pep-nanotrap (data not shown). Next, we tested whether the Pep-nanotrap is also suited to capture proteins displaying the PepTag at their C-terminus. Thus, we performed immunoprecipitation of the structural protein vimentin comprising a PepTag at its C-terminus (mCherry-VIM_Pep_) from the soluble protein fraction of transiently transfected HEK293T cells (**Figure 1 A**). The results show that Pep-tagged vimentin was efficiently precipitated by the Pep-nanotrap (**Figure 1 C**). In contrast to _Pep_GFP, we detected only minor amounts of the mcherry-VIM_Pep_ in the non-bound fraction. Additionally, western blot analysis revealed co-precipitation of endogenous vimentin along with the tagged version indicating an incorporation of mCherry-VIM_Pep_ into the network of the endogenous intermediary filaments. From these findings we conclude that the PepNb binds its epitope in the context of exogenously expressed proteins and can be used as an affinity matrix to capture recombinant Pep-tagged proteins. However, the position of the PepTag within the recombinant protein seems to influence the binding capacity of the Pep-nanotrap. Our observations suggest that the high binding affinity of the PepNb as shown previously ^29^ is valid only for the C-terminal position of the PepTag.

**Figure 1:**
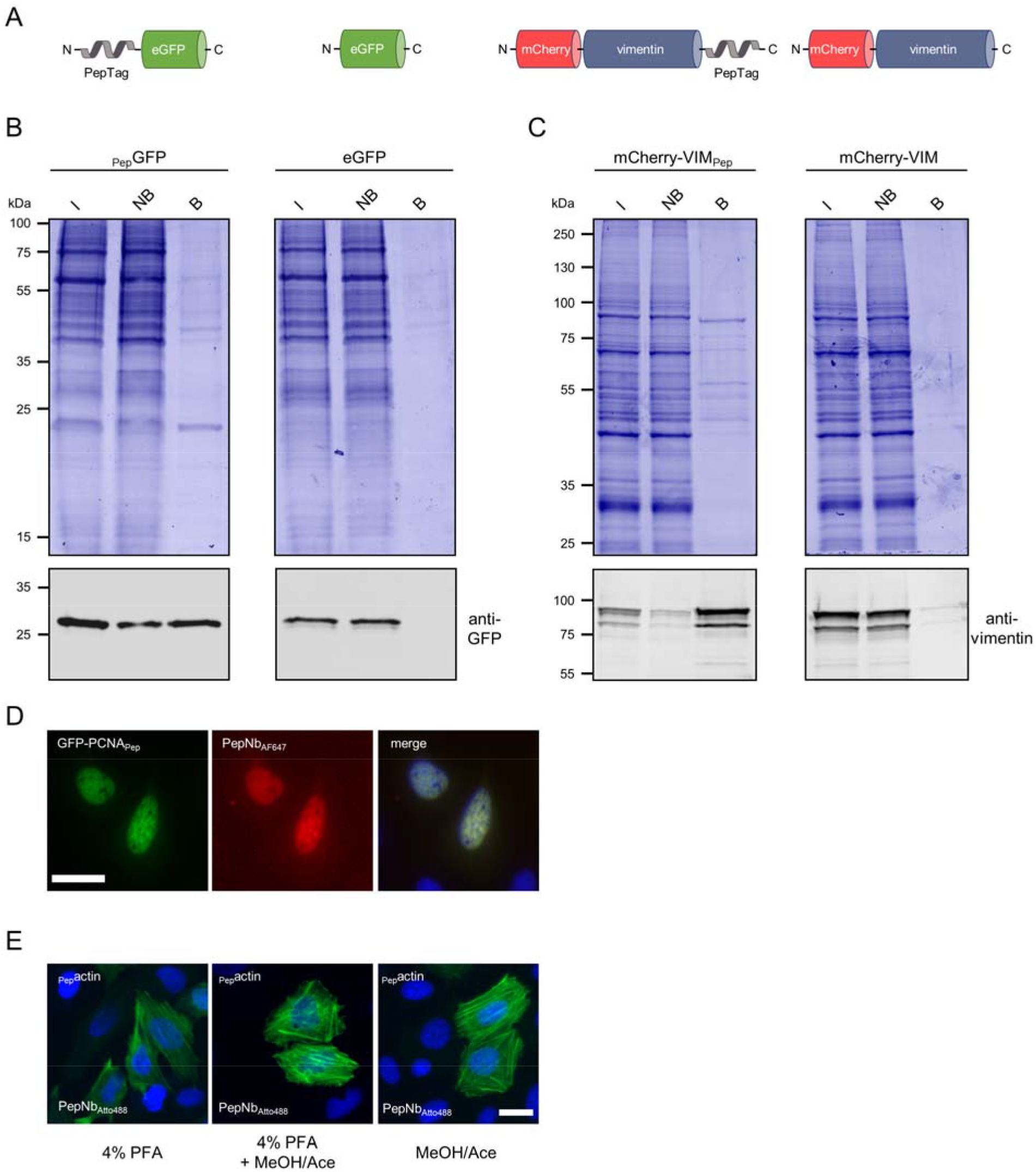
Purification and detection of Pep-tagged proteins with functionalized PepNb. (**A**) Schematic illustration of Pep-tagged fusion proteins and control proteins used for immunoprecipitation. (**B, C**) For immunoprecipitation soluble protein fractions of HEK293T cells either expressing GFP with an N-terminal PepTag (_Pep_GFP), eGFP, mCherry-vimentin comprising the PepTag at the C-terminus (mCherry-VIM_Pep_), or mCherry-vimentin (mCherry-VIM) were incubated with the Pep-Nb immobilized on sepharose-beads (Pep-nanotrap). Input (I), non-bound (NB) and bound (B) fractions were separated by SDS-PAGE and analyzed either by Coomassie Blue staining (top) or immunoblot with an anti-GFP antibody (bottom). (**D**) U2OS cells expressing GFP-PCNA_Pep_ were fixed with methanol/acetone and stained with PepNb conjugated to AlexaFluor647 (PepNbAF647, red) and nuclei were stained with DAPI (blue). Scale bar 25 μm. (**E**) U2OS cells expressing _Pep_actin were fixed using either 4% PFA, 4% PFA followed by methanol/acetone (MeOH/Ace) or MeOH/Ace alone and stained with PepNb conjugated to Atto488 (PepNbAtto488, green). Nuclei were stained with DAPI (blue). Scale bar 25 μm.

### Detection of Pep-tagged proteins by direct immunofluorescence

To test whether the PepTag/PepNb system is also suitable for direct detection of epitope-tagged proteins in immunofluorescence (IF), we coupled the organic dyes Alexa Fluor 647 (AF647) and Atto 488 site specifically to the C-terminus of the PepNb using Sortase A ^35^. First we monitored the performance of the PepNb in IF by fluorescence colocalization studies in U2OS cells transiently expressing a fusion construct encoding GFP-PCNA_Pep_, which localizes at discrete sites of DNA replication in the nucleus forming characteristic patterns during the S-phase of the cell cycle ^36^. IF staining with the PepNb_AF_647 revealed a clear co-localization of the Nb and the GFP signal at DNA replication foci in the nucleus (**Figure 1 D**). Secondly, we performed IF staining of actin filaments using the PepNb_ATTO488_. In this experiment we additionally tested the compatibility of the PepTag with standard fixation methods. Thus, we transiently expressed Pepactin in U2OS cells and fixed the cells 24 h post transfection using PFA PFA followed by methanol/acetone treatment or methanol/acetone alone. For all tested fixation methods IF staining with the PepNb_ATTO488_ showed characteristic fibers of the actin cytoskeleton (**Figure 1 E**). From these data we concluded that addition of the PepTag does not affect the characteristic localization of proteins and fluorophore-conjugated PepNbs are suitable to directly visualize antigens comprising this epitope tag in immunofluorescence staining independent of the applied fixation method.

### Detecting PepCB binding of Pep-tagged proteins in living cells

For intracellular applications we fused the PepNb to TagRFP or eGFP thereby generating Pep-chromobody (PepCB) expression constructs. In a first step we performed intracellular immunoprecipitations (IC-IPs) to test the antigen binding capability of the PepCB. Thus we co-transfected HEK293T cells with the PepCB (TagRFP) in combination with _Pep_GFP or eGFP as control and incubated the soluble protein fraction with the RFP-trap to pull down PepCB or the GFP-trap to pull down the antigen ^14,30^. Immunoblot analysis of the input, non-bound and bound fractions using anti-GFP or anti-TagRFP antibodies showed a clear enrichment of the PepCB in the bound fraction of _Pep_GFP. A lower amount of _Pep_GFP is also co-precipitated along with the PepCB (**Figure 2 A,** left panel). Notably, for co-expression of PepCB and eGFP no co-precipitation was observed (**Figure 2 A,** right panel). To further confirm intracellular chromobody binding we visualized the interaction of the PepCB with the Pep-tagged proteins in living cells using the Fluorescent Two-Hybrid Assay (F2H, ^8^). We performed triple transfections of BHK cells comprising a stably integrated lac operator (lacO) array ^37^ with constructs encoding for a fusion of lac inhibitor-GFP binding protein (lacI-GBP), _Pep_GFP or eGFP and the PepCB. Fluorescence imaging showed a clear localization of both eGFP constructs at the spot-like lacO array in the nucleus of BHK cells (**Figure 2 B**). A recruitment of the PepCB to the lacO-spot was only observable in the presence of _Pep_GFP (**Figure 2 B,** left panel) whereas upon co-expression of eGFP a dispersed distribution of the PepCB signal was observable (**Figure 2 B** right panel). To further validate these findings, we transfected U2OS cells with constructs encoding GBP-lamin B1 ^32^, _Pep_GFP or eGFP in combination with the PepCB. Fluorescence images of triple transfected cells revealed the recruitment of both GFP constructs to the nuclear lamina via the lamin-GBP construct. However, only in cells expressing _Pep_GFP the PepCB was detectable at the nuclear membrane (**Supplementary Fig. 1,** left panel). In summary, both the biochemical binding analysis and the fluorescence colocalization studies showed a specific intracellular binding capacity of the PepCB in living cells.

**Figure 2:**
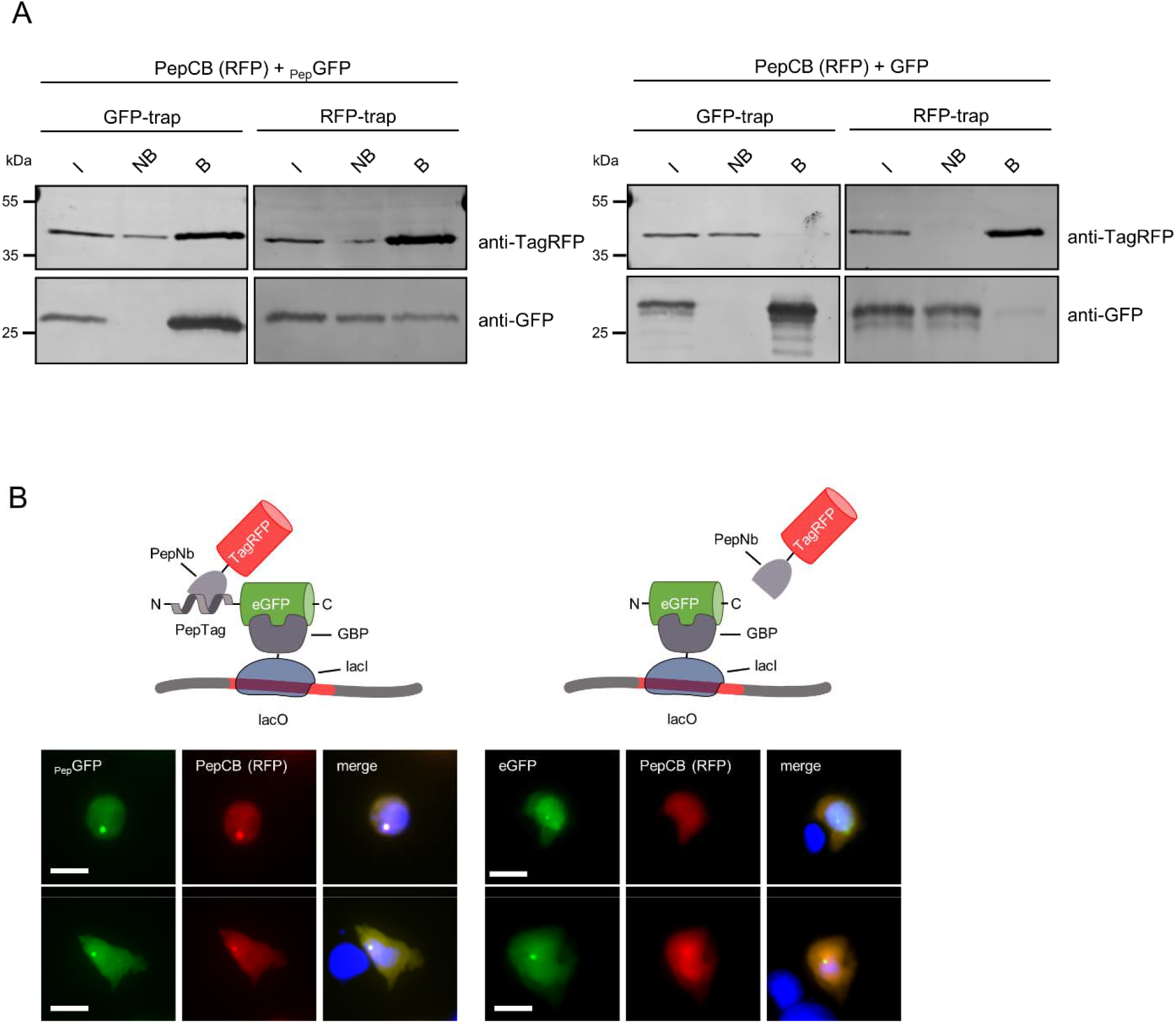
PepCB binds to its antigen in living cells. (**A**) Intracellular immunoprecipitation (IC-IP). Lysates of HEK293T cells expressing _Pep_GFP or GFP in combination with the PepCB comprising TagRFP as detectable moiety were subjected to immunoprecipitation with the GFP-trap or the RFP-trap. Input (I), non-bound (NB) and bound fractions (B) were analyzed by immunoblot with an anti-TagRFP antibody (upper panel) and an anti-GFP antibody (lower panel). (**B**) F2H assay to test the functionality of the PepCB *in cellulo*. Schematic drawing shortly outlines the assay principle (top). Transgenic BHK cells containing *lac* operator repeats (lacO) were co-transfected with lacI-GBP, PepCB (TagRFP) and _Pep_GFP (left panel) or eGFP (right panel). lacI-GBP binds to the lacO array and recruits GFP (bait protein, visible as a green spot, left column). PepCB (prey protein) binds anchored _Pep_GFP but not GFP (middle column). Nuclei of cells were counterstained with DAPI (right column). Representative images are shown. Scale bar 25 μm.

### Visualization of intracellular proteins using the PepCB

Next, we studied whether the PepTag/PepCB system is applicable to different cellular structures. Thus we transiently co-expressed the PepCB in combination with Pep-tagged components of microtubules (_Pep_GFP-tubulin), microfilaments (Pepactin), intermediary filaments (mCherry-vimentinPep) and the replication machinery (GFP-PCNA_Pep_) in U2OS cells and monitored antigen recognition of the PepCB by live-cell fluorescence imaging. All ectopically expressed proteins were efficiently incorporated in the corresponding cellular structures and could be imaged at their innate cellular localization (**Figure 3**). As shown for microtubules, PepCB visualized these structures exclusively in cells expressing _Pep_GFP-tubulin (**Figure 3 A,** left panel) but not in the absence of the PepTag (**Figure 3 A,** right panel). Moreover, incorporation of Pepactin into microfilaments could be visualized with the PepCB regardless of which fluorescent protein (eGFP or TagRFP) was used as detectable moiety for chromobody generation (**Figure 3 B**). To test whether the PepCB can also detect structural proteins comprising a C-terminal PepTag we performed live-cell imaging of U2OS cells expressing PepCB in combination with mCherry-VIM_Pep_ where we could detect a clear colocalization of the CB signal with the mCherry signal at cytoplasmic vimentin structures only in cells expressing the Pep-tagged version of mCherry-vimentin (**Figure 3 C,** right panel). Finally, to further confirm the binding capacity of the PepCB to C-terminally tagged proteins, we performed live cell imaging of U2OS cells expressing PepCB in combination with GFP-PCNA_Pep_. The images showed a clear colocalization of the PepCB signal and the GFP signal at DNA replication foci within the nuclei of double transfected cells (**Figure 3 D,** left panel), whereas we detected a diffuse PepCB distribution in cells coexpressing untagged GFP-PCNA (**Figure 3 D,** right panel). These data demonstrate the capability of the PepCB to visualize Pep-tagged antigens in different compartments in living cells. As tested for four different structural proteins neither the addition of the PepTag nor binding of the PepCB showed any sign of disturbing the characteristic localization of the antigen.

**Figure 3:**
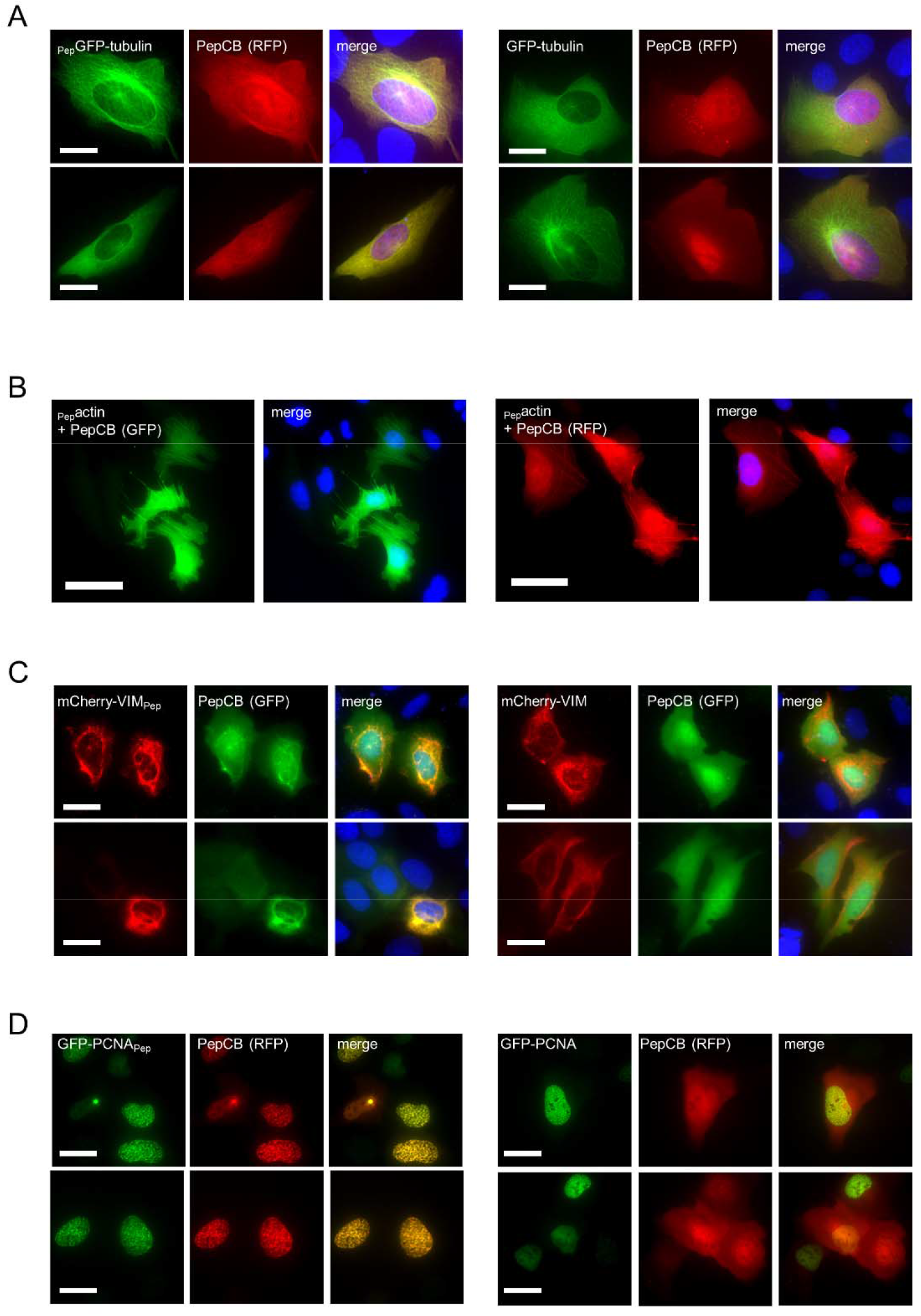
PepCB visualizes Pep-tagged proteins in living cells. Representative fluorescence images of living U2OS cells transiently expressing PepCB in combination with (**A**) _Pep_GFP-tubulin (left panel) or GFP-tubulin as control (right panel); (**B**) _Pep_actin, (**C**) mCherry-VIM_Pep_ (left panel) or mCherry-VIM as control (right panel); (**D**) GFP-PCNA_Pep_ (left panel) or GFP-PCNA (right panel). (A-D) Nuclei were stained with Hoechst33258. (A, D) Scale bar 25 μm, (B, C) Scale bar 50 μm.

### The PepCB binds only transiently to its epitope within living cells

Our findings suggest that the PepCB visualizes the Pep-tagged target structure without interfering with its localization or function. As demonstrated previously for other CBs this could be due to a transient binding mode ^9, 14, 33^ To analyze intracellular binding properties, we determined the turnover-rate of the PepCB upon binding to Pepactin by fluorescence recovery after photobleaching (FRAP). U2OS cells transiently expressing PepCB and _Pep_actin or respective control constructs (GFP-actin or actin-CB) were photobleached at indicated regions (**Figure 4 A**). Quantitative evaluation of the FRAP data revealed that only ~30% of the GFP-actin fluorescence signal recovered within 40 s (**Figure 4 B**) indicating a stable incorporation of fluorescently labeled GFP-actin in the cytoskeletal filaments. In contrast, both CBs targeting either endogenous actin (actin-CB) or _Pep_actin (PepCB) showed significantly higher recovery rates of ~90% within the analyzed time frame (**Figure 4 B**). Notably, the PepCB relocalized even faster to actin structures (t_1/2_ = 1.08 s) compared to the actin-CB (t_1/2_ = 2.01 s). Such fast recovery rates are characteristic for transient antigen binding and in accordance with previous findings ^33^. From these data we concluded that despite of high affinities determined for the original nanobody VHH 2E7 (K_D_: 0.6 nM) *in vitro* ^29^ the PepCB display high off-kinetics *in cellulo*. Considering published data on transient binding chromobodies it is conceivable that such a binding mode results in only minor influence on the function and dynamics of the target protein ^9,33^.

**Figure 4:**
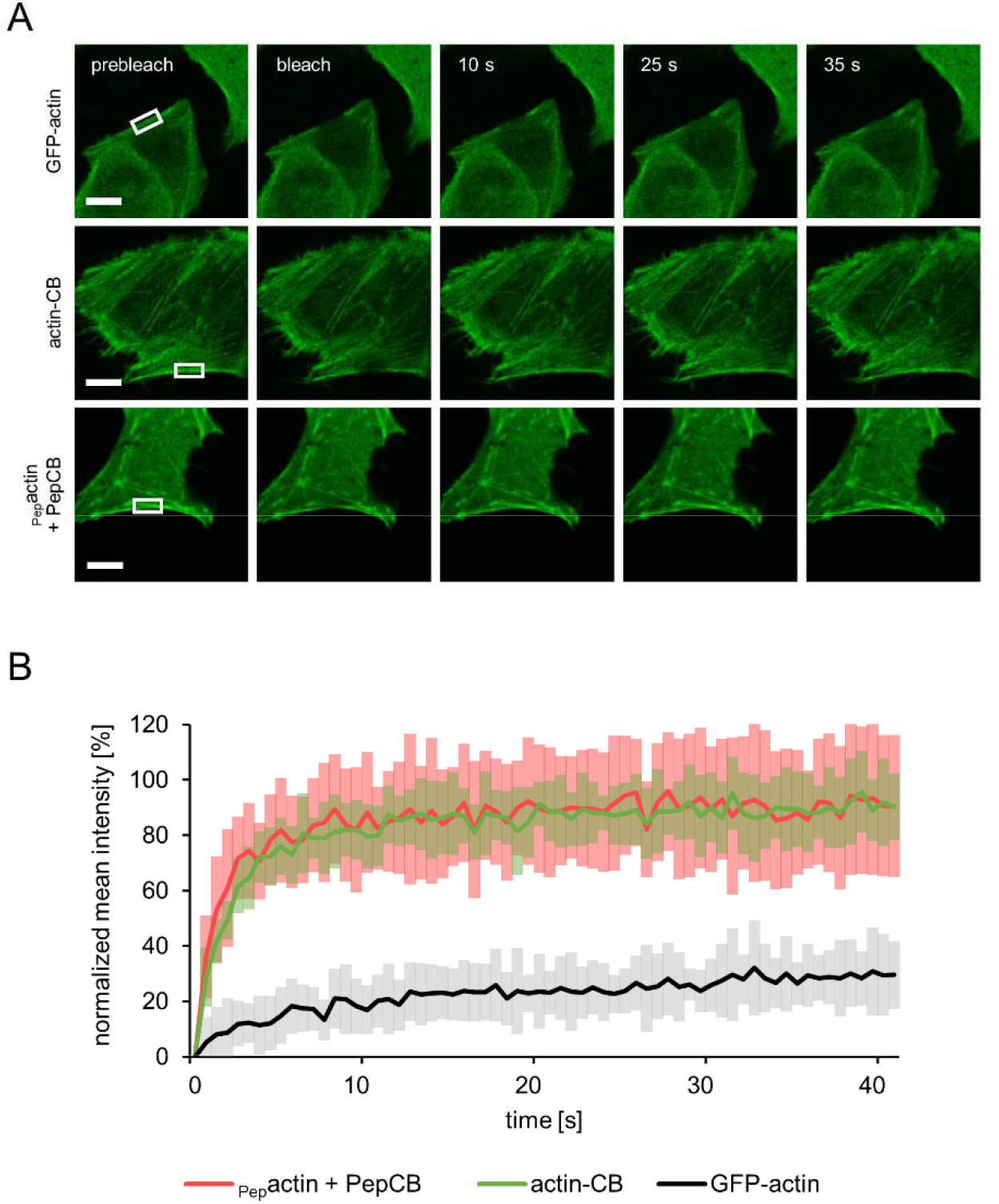
Quantification of intracellular binding properties of the PepCB. (**A**) Fluorescence recovery after photobleaching (FRAP) analysis of GFP-actin (upper row), actin-chromobody (actin-CB, middle row) and PepCB (eGFP) in combination with Pepactin (lower row) transiently expressed in U2OS cells. Shown are representative images at indicated time points before and after photobleaching of a defined region (white box). Scale bar 10 μm. (**B**) Quantification of FRAP data showing mean values of fluorescence signal in photobleached regions. Fluorescence was normalized to its intensity before bleaching. PepCB shows a recovery to 90.5 ± 25.6% with a halftime of 1.1 s. actin-CB recovered to 90.4 ± 12.1% with a halftime of 2.0 s whereas the recovery of GFP-actin amounted to 29.5 ± 12.2% with a half time of 20.3 s. n = 15, N = 1. Data are represented as mean ± stds.

### Intracellular visualization of protein dynamics using the PepCB

Next, we investigated the ability of the PepCB to trace dynamic changes of its antigen in living cells, again using GFP-PCNA_Pep_ and _Pep_actin. GFP-PCNA_Pep_ constitutes a special challenge to live-cell microscopy because it is an essential component of the replication machinery that concentrates at replication foci in S phase and shows a diffuse pattern in G1 and G2 ^36^. We followed the subcellular distribution of GFP-PCNA_Pep_ and the red fluorescent PepCB in non-synchronized cells throughout different stages of the cell cycle by taking images every 60 min for 12 h (**Figure 5 A, Supplementary Video 1**). At the beginning of the time series both GFP-PCNA_Pep_ and the PepCB showed a dispersed nuclear localization indicative for G1 phase shifted to more punctate structures which are characteristic for the progression of cells into S phase. Throughout the complete observation period the PepCB signal colocalizes with the GFP signal derived from its antigen GFP-PCNA_Pep_.

**Figure 5:**
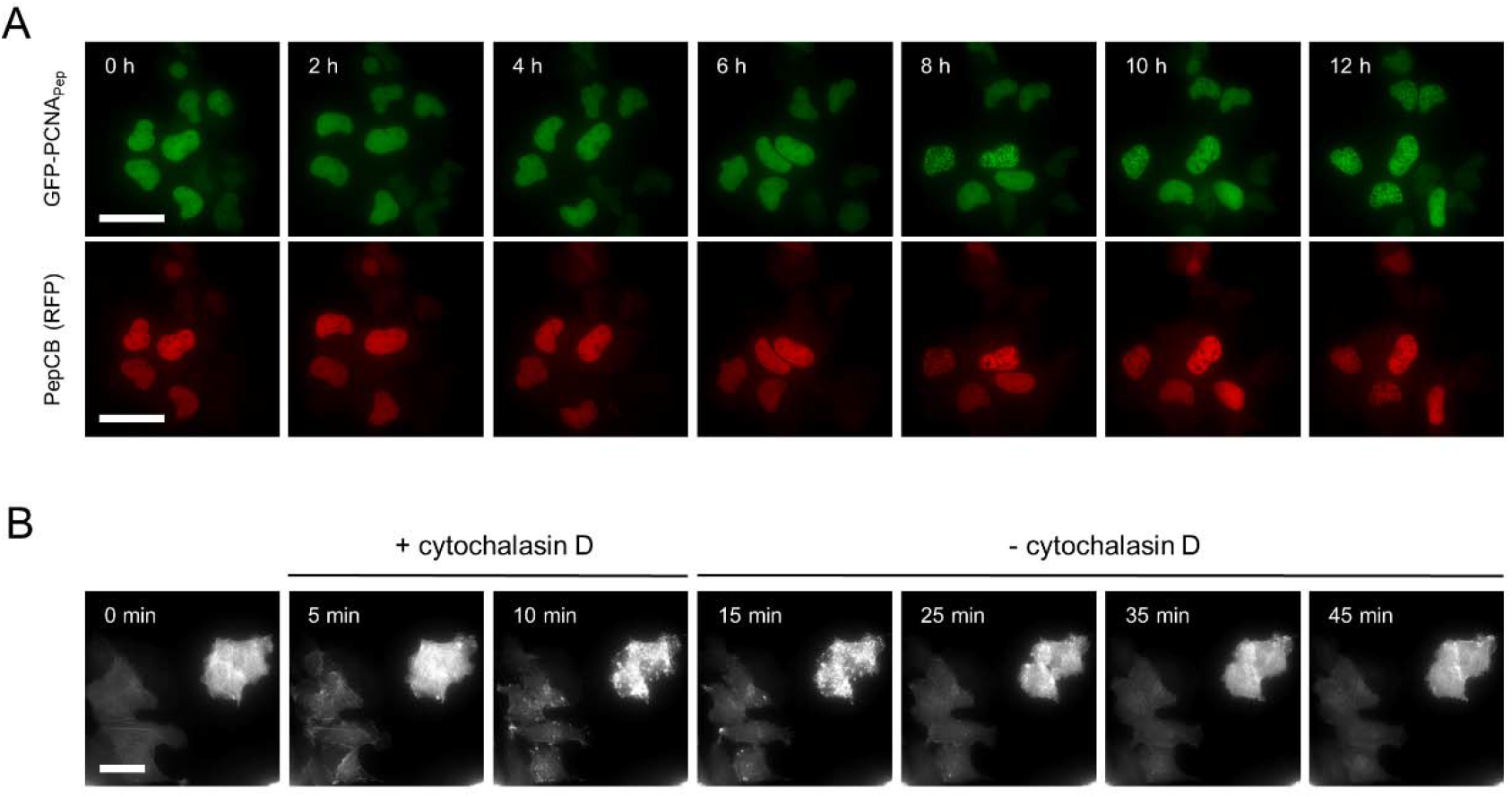
PepCB visualizes distribution and reorganization of Pep-tagged proteins in living cells. (**A**) Time-lapse microscopy of U2OS cells co-transfected with constructs encoding for PepCB (TagRFP) and GFP-PCNA_Pep_. Images were taken every hour, shown are representative images at indicated time points. Scale bar 50 μm. (**B**) U2OS cells expressing PepCB (TagRFP) and _Pep_actin. Immediately after starting the time series cells were treated with 2 μM cytochalasin D (actin polymerization inhibitor) for 10 min. Subsequently, cytochalasin D was removed and cells were continuously imaged for additional 30 min. Shown are representative fluorescence images displaying chromobody signal at indicated time points. Scale bar 50 μm.

To test whether the PepCB also visualizes dynamic changes of its antigen upon compound treatment, we performed time-lapse imaging on U2OS cells co-expressing the PepCB and _Pep_actin which were initially treated with the actin-modulating compound cytochalasin D, for 10 min. Subsequently, we removed the compound and continuously imaged the cells for 30 min (**Figure 5 B, Supplementary Video 2**). The PepCB signal revealed a rapid re-organization of filamentous actin within 5 min of cytochalasin D treatment followed by the reappearance of actin fibers upon removal of the compound within 30 min. These time-lapse analyses showed that the PepCB can trace essential components of the replication machinery as well as tightly organized cytoplasmic structures without major impact on cell viability.

### Antigen-mediated stabilization of the PepCB

Recently, we demonstrated that intracellular CB levels respond to the amount of their respective antigens, a phenomenon termed antigen-mediated chromobody stabilization (AMCBS) ^30^. Using turnover-accelerated CBs displaying an arginine as N-terminal amino acid residue (Ub-R-CBs) we employ AMCBS to visualize and monitor fast reversible changes in cellular antigen concentration upon compound treatment by quantitative live-cell imaging ^30^. Additionally, we established a protocol for site-directed integration of CB encoding sequences into the genomic safe-harbor AAVS-1 locus of human cells using the CRISPR/Cas9 technology to generate stable cell lines displaying an homogenous CB expression ^34^. Here we applied this approach to generate U2OS cells stably expressing a turnover-accelerated version of the PepCB (Ub-R-PepCB) and analyzed its performance to visualize and quantify changes of Pep-tagged antigen levels by quantitative live-cell imaging.

Starting from a polyclonal pool of U2OS cells expressing the Ub-R-PepCB from the safe harbor locus we performed monoclonal selection and PCR-based genotyping to identify and validate positive clones. For U2OS clone E02 our data revealed an amplicon at the size of ~1018 bps indicating a correct integration of the CB transgene into the AAVS1 locus which was further confirmed by a single amplicon at the size of ~4346 bps, suggesting a homologous integration of Ub-R-PepCB into the AAVS1 locus of the monoclonal U2OS cells (**Supplementary Figure 2**).

To investigate whether the Ub-R-PepCB is stabilized in the presence of its antigen, we transiently transfected CRISPR-engineered U2OS_E02 cells with _Pep_GFP, _Pep_GFP-NLS and GFP-PCNA_Pep_. By comparing nuclear-localized _Pep_GFP and homogenous distributed _Pep_GFP we additionally aimed to analyze the influence of these two different cellular compartments on AMCBS. For quantitative analysis we determined the average CB fluorescence signal within transfected cells in comparison to cells expressing the corresponding, non-tagged constructs. Notably, in these cells we hardly detect any PepCB signal in line with a fast turnover of the stably expressed Ub-R-PepCB (**Figure 6**).

**Figure 6:**
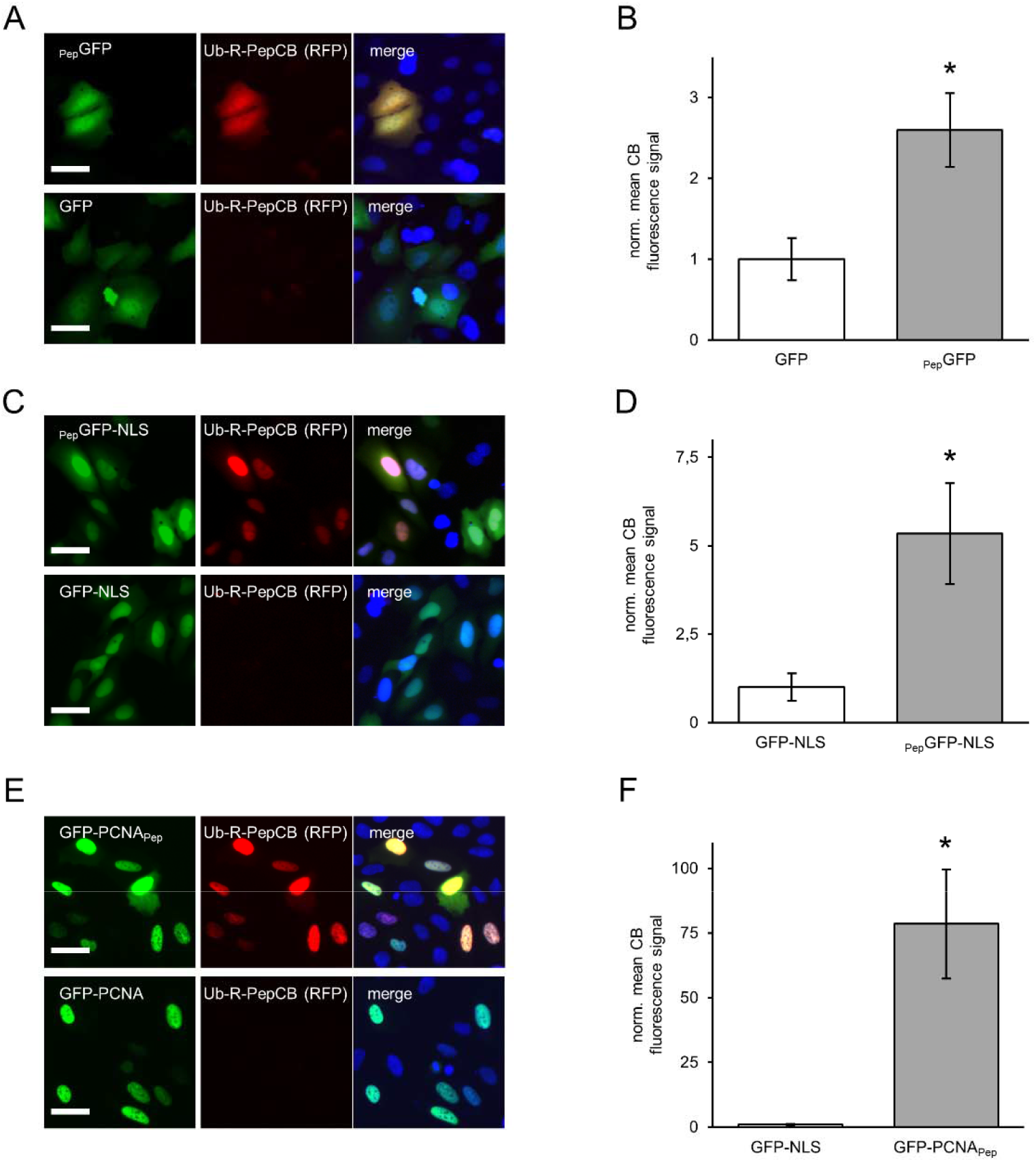
PepCB is stabilized in the presence of Pep-tagged antigens. (**A**) Fluorescence images of living U2OS E02 cells comprising a homologues integration of the turnover-accelerated PepCB (Ub_R_PepCB) at the AAVS-1 locus upon transfection with _Pep_GFP (top) or eGFP as control (bottom). (**B**) Bar chart displaying normalized mean nuclear chromobody fluorescence detected in eGFP or _Pep_GFP expressing U2OS_E02 cells. Mean fluorescence was calculated from three samples (n=3; > 200 cells) and normalized to GFP-transfected cells (set to 1). (**C**) Fluorescence images of cells as described in (**A**) upon expression of _Pep_GFP-NLS (top) or GFP-NLS as control (bottom). (**D**) Bar chart displaying the normalized mean chromobody fluorescence detected in nuclei of GFP-NLS or _Pep_GFP-NLS expressing U2OS_E02 cells (n=3; > 200 cells). Mean fluorescence was calculated from three samples (n=3; > 200 cells) and normalized to GFP-NLS expressing cells (set to 1). (**E**) Fluorescence images of cells as described in (**A**) upon expression of GFP-PCNA_Pep_ (top) or GFP-PCNA (bottom). (**F**) Bar chart of normalized mean nuclear fluorescence of PepCB detected in GFP-PCNA or GFP-PCNA_Pep_ expressing cells. Mean fluorescence was calculated from three samples (n=3; > 200 cells) and normalized to GFP-PCNA expressing cells (set to 1). (**A, C, E**) Shown are cells 24 h post transfection. For nuclear segmentation, cells were stained with Hoechst33258. Scale bar: 50 μm. (**B, D, F**) Data are represented as mean ± stds. For statistical analysis student’s t-test was performed, * p < 0.05.

Upon transient transfection of _Pep_GFP we observed a stabilization of the UB-R-PepCB exclusively in cells expressing the antigen (**Figure 6 A**). Quantification of Ub-R-PepCB signal revealed a stabilization factor of ~2.6 (**Figure 6 B**). Cells expressing the nuclear localized _Pep_GFP-NLS showed a stronger Ub-R-PepCB signal with a calculated stabilization factor of ~5.3 (**Figure 6 C, D**). From these findings we concluded that antigen binding within the nucleus increases Ub-R-PepCB stabilization substantially. This becomes even more obvious when we transiently transfected nuclear localized GFP-PCNA_Pep_ (**Figure 6 E**). Deduced from the observable GFP signal this construct showed substantially higher expression rates compared to _Pep_GFP or _Pep_GFP-NLS. In parallel we observed a significant brighter Ub-R-PepCB signal (**Figure 6 E**) and calculated a stabilization factor of ~78 (**Figure 6 F**).

In summary our data demonstrated that the stably expressed, turnover-accelerated Ub-R-PepCB is stabilized in the presence of various Pep-tagged antigens. This is in accordance to previous findings of antigen-mediated stabilization of intrabodies and chromobodies targeting endogenous epitopes ^30,39–41^. In addition to our previous data we showed that the stabilization effect is much stronger for antigens localized within the nuclear component compared to cytoplasmic localized ones. Although the precise molecular and structural mechanisms for this phenomenon remain to be elucidated we hypothesized that the antigen-bound PepCB which is degraded via the ubiquitin proteasomal system has restricted access to the proteasome localized in the cytoplasm.

## Summary and Outlook

Epitope tagging is not only an effective way to facilitate expression and purification of recombinant proteins but also a favored approach to study biogenesis, localization and molecular interactions of proteins of interest (PoIs) when lacking well-defined and reproducible capture and/or detection reagent. In this context we and others have developed nanobodies which on multiple occasions have proven to be valuable and versatile tools for applications ranging from one-step purification to super-resolved imaging of proteins comprising a small peptide tag ^26–28,35^. Here we build on previous work describing a nanobody in complex with a short helical peptide within the gp41 protein of HIV ^29^ and designed numerous mammalian expression constructs displaying the minimal (12 AA) epitope (PepTag) either on the C- or N-terminus. We demonstrated the capability of functionalized versions of the nanobody (PepNb) for immunoprecipitation (IP) and direct immunofluorescence (IF) of Pep-tagged proteins. Focusing on live-cell imaging we generated intracellularly functional PepCB constructs and showed that these PepCBs efficiently target and trace Pep-tagged antigens in living cells. Notably, neither the short PepTag nor binding to the PepCB affects localization and dynamics of Pep-tagged proteins analyzed in different cellular compartments which is likely due to a high-off rate as indicated from fast recovery rates measured by FRAP. Finally, following up our recent findings on antigen-mediated chromobody stabilization ^30,34^ we showed for the first time that a turnover-accelerated PepCB can be used as a broadly applicable generic biosensor to monitor changes in cellular concentrations of antigens harboring a small and inert peptide tag simply by quantitative live-cell imaging. In summary the PepTag/PepCB tagging system described herein provides a highly versatile and broadly applicable approach to study PoIs in different experimental settings.

## Abbreviations

AAVS1: adeno-associated virus integration site 1
ACT-CB: actin-chromobody
AMCBS: antigen-mediated chromobody stabilization
CB: chromobody
CRISPR: clustered regularly interspaced short palindromic repeats
DAPI: 4’,6-Diamidin-2-phenylindol
DMEM: Dulbecco’s Modified Eagle Medium
EDTA: ethylenediaminetetraacetic acid
F2H: Fluorescent Two-Hybrid Assay
FCS: fetal calf serum
FRAP: fluorescence recovery after photo-bleaching
GBP: GFP binding protein
IF: immunofluorescence
IMAC: immobilized metal affinity chromatography
IP: immunoprecipitation
Nb: nanobody
PAGE: polyacrylamide gel electrophoresis
PCNA: proliferating cell nuclear antigen
PFA: paraformaldehyde
PEI: Polyethylenimine
PMSF: phenylmethylsulfonyl fluoride
SDS: sodium dodecyl sulfate
Ub: Ubiquitin
VIM: vimentin

## Author contributions

B.T. and U.R. conceived the study. B.T., S.S., P.D.K. and U.R. performed all experiments. B.T., S.S. and U.R. analyzed the data. B.T., S.S. and U.R. wrote the manuscript

## Acknowledgements

The authors gratefully acknowledge the Ministry of Science, Research and Arts of Baden-Württemberg (V.1.4.-H3-1403-74) for financial support.

