## Supplementary Information for "A novel epitope tagging system to visualize and monitor antigens in live cells with chromobodies"

| oligonuclotide name | sequence (5‘ – 3‘) |
| --- | --- |
| PepTag-ACTB-for | ATAGGTACCTGAAAGATCAGCAGCTGCTGGGCATTTGGGGAGGCAGCGATGATGATATCGCCGCGCTCGTC |
| PepTag-ACTB-rev | TATGGTACCTTTCCACCGCCATGGTGGCGGTGGCGACCGGTAGCGCTAG |
| AAVS1-HA-L-for | CCTCTCTAGTCTGTGCTAGCTC |
| AAVS1-HA-R-rev | GAAGGAGGAGGCCTAAGGATGG |
| AAVS1-vor-HA-L-for | CGGAACTCTGCCCTCTAACG |
| AAVS1-T2A-rev | GGGATTCTCCTCCACGTCAC |
| PCNA-PepTag-for | CTGAAGGACCAGCAGCTCCTCGGCATCTGGTAGTCTAGAGTCGAGATC |
| PCNA-PepTag-rev | GTACCTCTCCACGGCGGATCCCCCGCCTCCAGATCCTTCTTCATCCTC |
| PepNB-DNA-fragment | AGATCTCCGGCCATGGCTGACGTGCAGCTGCAGGAGAGCGGCGGCGGCCTGGTGCAGCCCGGCGGCAGCCTGAGGCTGAGCTGCGCCGCCAGCGGCAACATCGTGAGCATCGACGCCGCCGGCTGGTTCAGGCAGGCCCCCGGCAAGCAGAGGGAGCCCGTGGCCACCATCCTGACCGGCGGCGCCACCAACTACGCCGACAGCGTGAAGGGCAGGTTCACCATCAGCAGGGACAACGCCAAGAACACCGTGTACCTGCAGATGAACAGCCTGAAGCCCGAGGACACCGCCGTGTACTACTGCTACGCCCCCATGATCTACTACGGCGGCAGGTACAGCGACTACTGGGGCCAGGGCACCCAGGTCACC |
| vimentin-PepTag-for | GAGGTACCTGAAGGACCAGCAGCTcCTcGGCATCTGGtgagatccaccggatctagataactgat |
| vimentin-PepTag-rev | CAGGTACCTCTCCACGGCGGATCCCCCGCCTCCtggttcaaggtcatcgtgatg |
| PepTag-eGFP-for | aaaggtacctgaaagatcagcagctgctgggcatttggggaggcagcGTGAGCAAGGGCGAGGAGC |
| PepTag-eGFP-rev | TTTCAGGTACCTTTCCACCGCCATGCTAGCGGATCTGACGGTTC |
| eGFP-NLS-for | AGAAGAGGAAGGTTTGATAAagcggccgcgactct |
| eGFP-NLS-rev | TCTTAGGGCTGCCTCCCTTGTACAGCTCGTCCATGCC |

Table S1: List of DNA oligonucleotids and synthesized gene fragments used in this study

|  | **Expression constructs/stable cell lines** |
| --- | --- |
| 1. | U2OS_E02 (this study) |
| 2. | BHK cells containing multiple lac-operon repeats ^1^ |
| 3. | PepCB (tagRFP) (this study) |
| 4. | PepCB (eGFP) (this study) |
| 5. | lamin-CB ^2^ |
| 6. | PCNA-CB-TagRFP ^3^ |
| 7. | PepNb-Myc-KKK-His_6_ (this study) |
| 8. | PepNb-Sort-His_6_ (this study) |
| 9. | AAVS1_EF1-α-Ub-R-ACT-CB ^4^ |
| 10. | AAVS1_EF1-α-Ub-R-PepCB (this study) |
| 11. | mCherry-vimentin ^5^ |
| 12. | mCherry-vimentin_Pep_ (this study) |
| 13. | BC2T-eGFP ^6^ |
| 14. | _Pep_GFP (this study) |
| 15. | _Pep_actin (this study) |
| 16. | BC2T-actin ^6^ |
| 17. | _Pep_GFP-tubulin (this study) |
| 18. | PamCherry-tubulin ^7^ |
| 19. | GFP-PCNA_Pep_ (this study) |
| 20. | GFP-PCNA ^8^ |
| 21. | eGFP-NLS (this study) |
| 22. | _Pep_GFP-NLS (this study) |
| 23. | lacI-GBP ^9^ |
| 24. | Lamin-GBP ^9^ |
| 25. | GFP-actin (Clontech) |
| 26. | actin-CB ^3^ |
| 27. | eGFP (Clontech) |

Table S2: List of expression constructs and stable cell lines used in this study

**Supplementary Figure 1:**

Nuclear lamina-based interaction assay of the PepCB and _Pep_GFP *in cellulo*.

Top panels illustrate assay principle: nuclear lamina-resident GBP-lamin B1 recruits co-expressed _Pep_GFP along with PepCB (TagRFP); control situation depicted on the right.

Representative fluorescence images of living U2OS cells transiently co-expressing PepCB, GBP-lamin B1, and _Pep_GFP (lower left panel) or eGFP (lower right panel). Fluorescence colocalization at the nuclear lamina indicates PepCB binding to _Pep_GFP. Scale bar 25 µm.

**Supplementary Figure 2:**

PCR-based genotyping of CRISPR-engineered U2OS cell line.

Schematic outline of genetic elements at the AAVS1 locus after integration of the PepCB. Arrows indicate the position of primers used to confirm correct genomic integration.

**Supplementary Video 1:**

U2OS cells transiently co-expressing GFP-PCNA_Pep_ and the red fluorescent PepCB. Time interval 1 h, scale bar 50 µm.

**Supplementary Video 2:**

U2OS cells transiently co-expressing _Pep_actin and the red fluorescent PepCB. Cell were exposed to cytochalasin D for 10 min, followed by 30 min recovery. Time interval 5 min, scale bar 50 µm.

**Supplementary Figure 1**

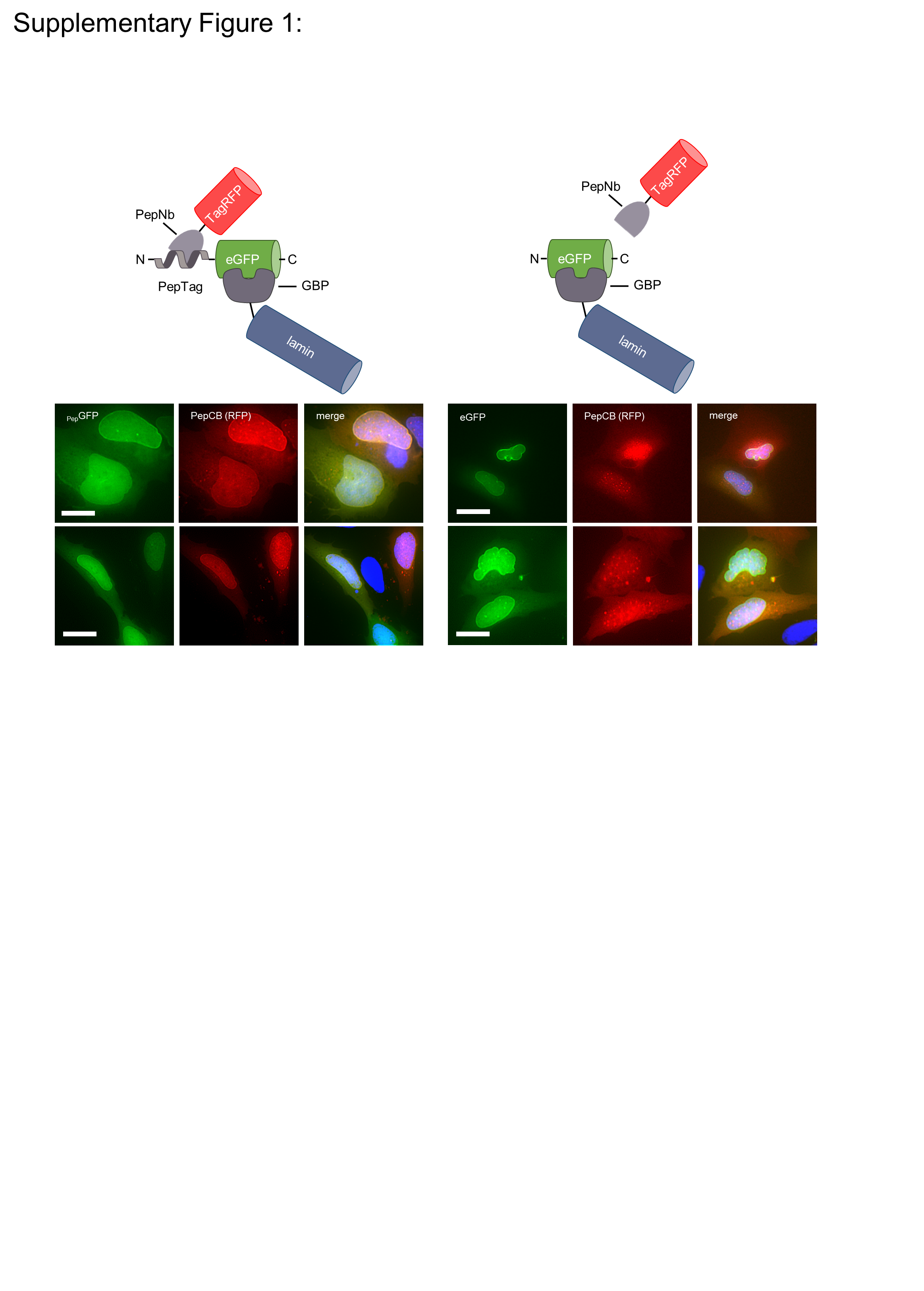

**Supplementary Figure 2**

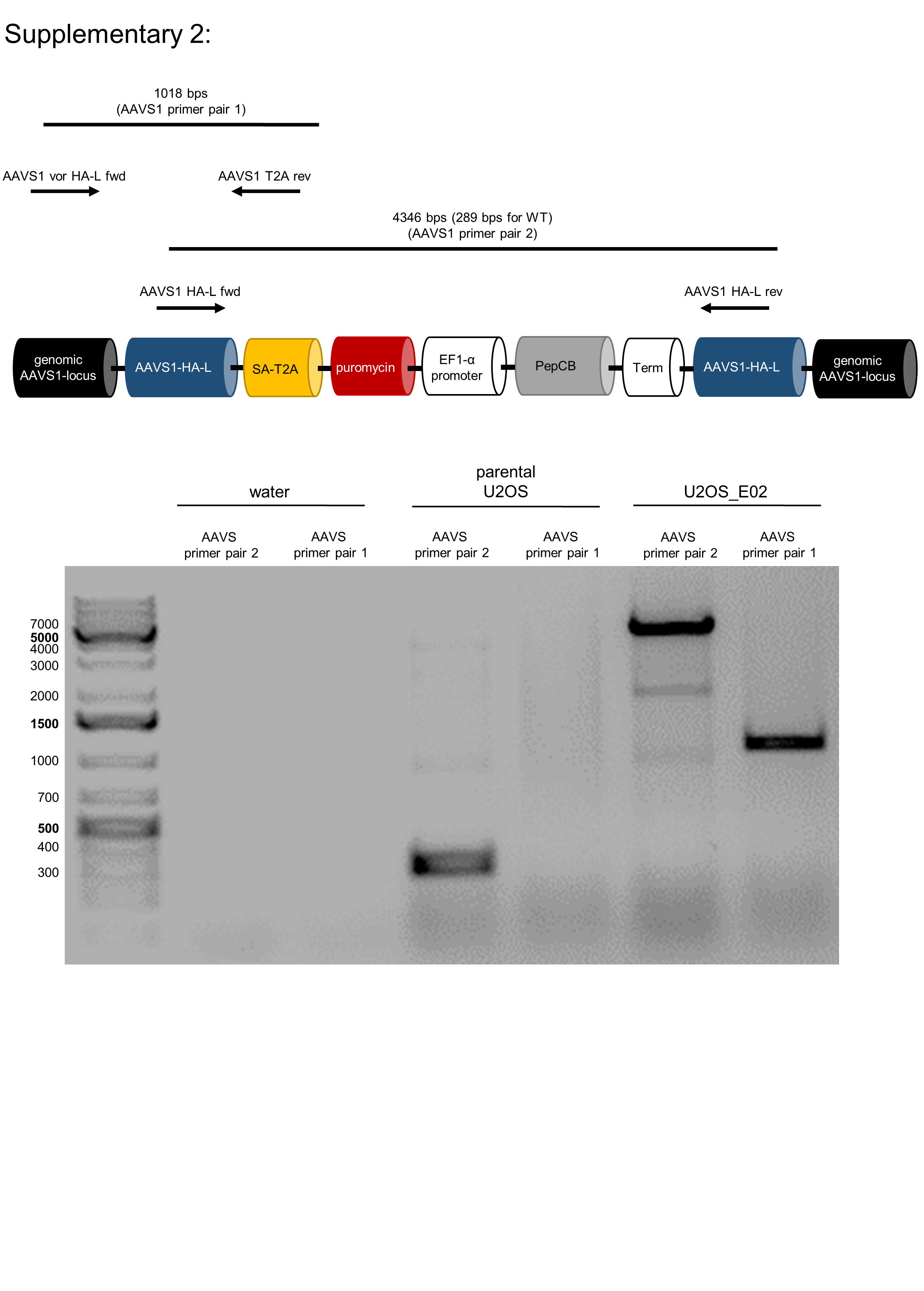
