## Supplementary figures and images for "A novel epitope tagging system to visualize and monitor antigens in live cells with chromobodies"

### Supplementary Video 1

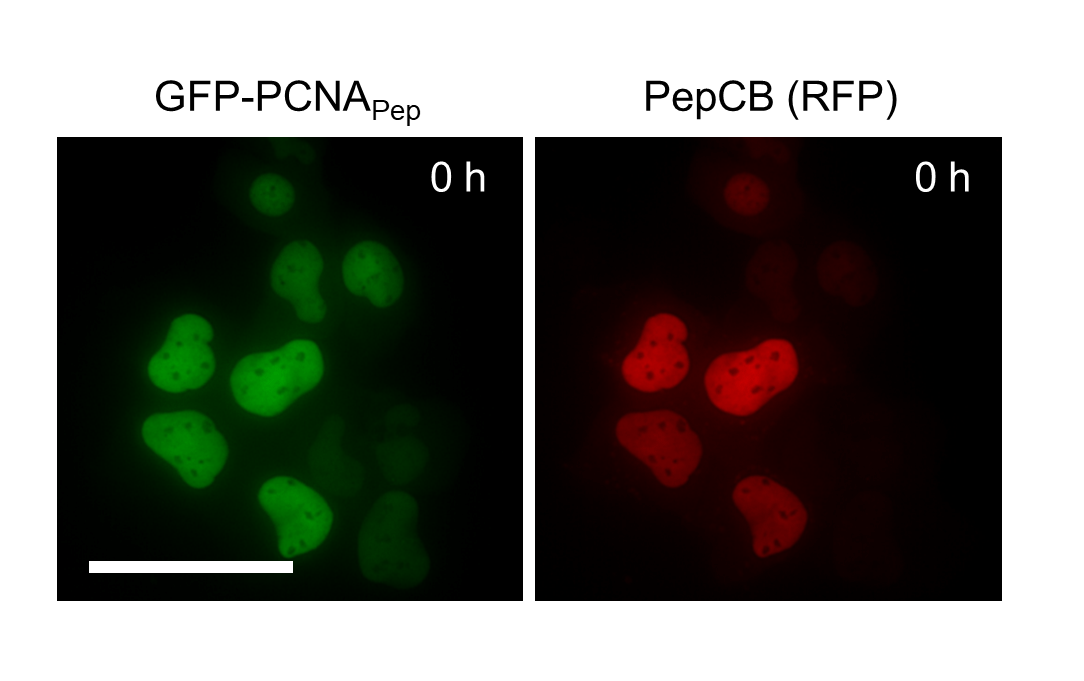
